# Direct visualization of epithelial microvilli biogenesis

**DOI:** 10.1101/2020.10.21.341248

**Authors:** Isabella M. Gaeta, Leslie M. Meenderink, Meagan M. Postema, Caroline S. Cencer, Matthew J. Tyska

## Abstract

Microvilli are actin bundle supported surface protrusions that play essential roles in diverse epithelial cell functions. To develop our understanding of microvilli biogenesis, we used live imaging to directly visualize protrusion growth at early stages of epithelial differentiation. Time-lapse data revealed that an “initiation complex” enriched in EPS8 and IRTKS appears at future sites of microvillus growth minutes before core actin bundle assembly. Elongation of a new core bundle occurs in parallel with the arrival of EZRIN and plasma membrane encapsulation. In addition to *de novo* growth, we also observed that new microvilli emerge from pre-existing protrusions. Additionally, we found that new microvilli can also collapse, characterized first by loss of membrane wrapping and Ezrin enrichment, followed by a sharp decrease in distal tip EPS8 and IRTKS. These studies are the first to offer a temporally resolved microvillus growth mechanism and highlight critical factors that drive this process.

## INTRODUCTION

Actin bundle-supported surface protrusions are one of the defining features of sophisticated eukaryotes, conferring cells with the ability to move and sense the surrounding environment. Examples include filopodia and microvilli, fingerlike protrusions that extend from the leading edge of crawling cells and the apical surface of polarized epithelial cells, respectively. In an evolutionary context, filopodia-like structures predate animal cells (Adl et al., 2012), whereas the evolution of microvilli occurred later, emerging in animal cells and choanoflagellates (Karpov and Leadbeater, 1998; Peña et al., 2016). In higher eukaryotes, these protrusive structures are found on the surface of diverse epithelial tissues, where they perform a variety of functions. In the intestinal tract and renal proximal tubule, microvilli comprise the densely packed brush border, which increases apical surface area for nutrient and filtrate absorption, respectively (Coudrier et al., 1988; Helander and Fandriks, 2014). Additionally, in the sensory epithelium, exaggerated microvilli referred to as stereocilia respond to mechanosensory stimuli that allow for hearing (Schwander et al., 2010). Despite their essential roles in a broad range of critical epithelial tissue functions, little is known about the temporal and mechanistic details that govern when or how microvilli grow from the apical surface. Here, we sought to identify the molecular events that govern the earliest steps of microvilli growth.

Microvilli ultrastructure is well characterized with data from transmission electron microscopy (TEM) and biochemical studies spanning nearly six decades. Moreover, proteomic analyses have revealed a comprehensive list of proteins resident in microvilli, many of which interact directly with the actin cytoskeleton (McConnell et al., 2011; Revenu et al., 2012). In transporting epithelia, microvilli extend microns from the cell surface supported by a core of ~30-40 actin filaments (Mooseker and Tilney, 1975) that are bundled in parallel by VILLIN, FIMBRIN (also known as PLASTIN-1), and ESPIN (Bartles et al., 1998; Bretscher and Weber, 1979b; Bretscher and Weber, 1980). This core actin bundle is also physically linked to the encapsulating plasma membrane by membrane-cytoskeleton linkers including EZRIN, MYO1A, and MYO6 (Berryman et al., 1993; Bretscher, 1983; Hegan et al., 2012; Howe and Mooseker, 1983) which mechanically stabilize the resulting protrusion. Within core bundles, actin filament barbed-ends, which represent favorable sites for new monomer incorporation, are oriented towards the distal tips (Mooseker and Tilney, 1975). Classic TEM studies also revealed that barbed-ends are embedded in an electron dense cap, referred to herein as the “distal tip complex”, which is predicted to control the growth of microvilli (Tilney and Cardell, 1970). Electron dense distal tip complexes are also found at the tips of filopodia and stereocilia (Rzadzinska et al., 2004; Svitkina et al., 2003) suggesting a universal function in protrusion formation.

Previous studies identified several factors that are enriched at the distal tips and may reside in the electron dense complex. For example, filopodia, microvilli, and stereocilia all contain MyTH4-FERM myosins, which deliver cargo and promote protrusion elongation; MYO10 is enriched at filopodial tips (Berg and Cheney, 2002), whereas MYO15 (Belyantseva et al., 2003), MYO3A (Salles et al., 2009) and MYO7A (Grati and Kachar, 2011) accumulate at stereocilia tips and the upper tip link densities, and MYO7B exhibits robust tip targeting in microvilli (Weck et al., 2016). In both stereocilia and microvilli, transmembrane adhesion molecules are delivered to the tip compartment by these unconventional myosins, to form tip-linking complexes that physically connect adjacent protrusions. In stereocilia, CDH23 and PCDH15 form *trans* heterophilic adhesion complexes (Kazmierczak et al., 2007), whereas microvilli are linked by a complex composed of CDHR2 and CHDR5 (Crawley et al., 2014b). Tip enriched adhesion factors in filopodia serve to link the tips of filopodia to the underlying substrate, including integrins and integrin regulatory/adaptor proteins such as Talins, Kindlin-2 and ICAP-1 (Jacquemet et al., 2019; Lagarrigue et al., 2015).

Although a great deal of evidence indicates that myosin motors and tip-linking adhesion complexes function in the organization of finger-like surface features, factors that control the initiation and elongation of protrusions remain less clear. In filopodia, molecules that promote elongation include ENA/VASP family proteins and formins, which likely function by antagonizing Capping Protein and/or by promoting direct addition of new profilin-actin monomers to filament barbed-ends (Bear et al., 2002; Breitsprecher and Goode, 2013; Ferron et al., 2007). In stereocilia, a tip enriched complex containing WHIRLIN, epidermal growth factor pathway substrate 8 (EPS8), and MYO15A promote elongation (Manor et al., 2011b). Additionally, the barbed end binder Capping Protein drives widening of stereocilia, and this function may be aided by its interaction with TWINFILIN-2 (Avenarius et al., 2017). The EPS8 family member EPS8L2 is also enriched at the distal tips of stereocilia, and loss of this protein affects stereocilia length and results in gradual deafness (Furness et al., 2013). In microvilli, tip enriched actin binding proteins EPS8 and the I-BAR protein insulin receptor tyrosine kinase substrate (IRTKS, also known as BAIAP2L1) promote protrusion elongation (Postema et al., 2018a). Additionally, EPS8L1a localizes to the distal tips of microvilli and interacts with EZRIN, and co-expression of EZRIN with either EPS8L1a or EPS8 disrupts microvillus morphology (Zwaenepoel et al., 2012). Finally, EPS8 exhibits robust enrichment the distal tips of filopodia (Chou et al., 2014).

Due to its exquisitely specific and robust enrichment at the distal tips of microvilli, stereocilia and filopodia, EPS8 is of particular interest as a factor that may initiate and/or drive protrusion growth. Indeed, studies in mouse models showed that knockout of EPS8 leads to a significant shortening of both intestinal microvilli and stereocilia (Postema et al., 2018a; Tocchetti et al., 2010; Zampini et al., 2011), suggesting EPS8 is involved in the elongation of these structures. EPS8 is also required for proper apical morphogenesis in the *C. elegans* intestine, where genetic ablation leads to gross defects in tissue morphology and lethality if both isoforms are lost (Croce et al., 2004). Consistent with these findings, our lab recently determined that EPS8 and IRTKS work together to promote microvilli elongation (Postema et al., 2018a) and promote directional persistence of motile microvilli early in differentiation, in a manner dependent on their actin binding activities (Meenderink et al., 2019). Mechanistically, EPS8 has been assigned both actin capping and bundling functions (Disanza et al., 2004; Hertzog et al., 2010). While its localization at the tips of protrusions is consistent with a potential role in capping barbed-ends, this function is at odds with shortened protrusions that result from EPS8 loss-of-function (Postema et al., 2018b; Tocchetti et al., 2010; Zampini et al., 2011). Additionally, tip enrichment is not entirely consistent with a role in filament bundling, as canonical parallel actin bundling proteins localize along the length of protrusions (Bartles et al., 1998; Bretscher and Weber, 1979a).

Though our understanding of microvillus ultrastructure and resident proteins has expanded in recent years, little is known about how protrusion growth is initiated at the cell surface during epithelial differentiation. This knowledge gap is at least in part due to the technical demands associated with making time-resolved observations of protrusion growth events. For example, microvilli are small, with a width below the diffraction limit (~100nm), and the signal from any fluorescent probe at the earliest stages of growth will necessarily be dim. Microvilli also typically extend from the surface in large numbers, making it difficult to visually isolate individual growth events. Moreover, our recent studies demonstrate that early in differentiation, microvilli are highly dynamic and are capable of translocating across the apical surface (Meenderink et al., 2019). Although small, dim, and crowded moving targets present a significant live cell microscopy challenge, addressing these hurdles would allow us to generate insight on factors involved in microvilli growth as well as the timing of their recruitment to the apical surface.

To this end, we developed a live imaging assay that enabled us to directly observe the *de novo* growth of individual microvilli on the surface of epithelial cells. Based on its ubiquitous appearance at the tips of microvilli in diverse epithelial tissues and cell culture models, we tested the idea that EPS8 represents a critical factor involved in the *de novo* growth of microvilli. Consistent with this hypothesis, we observed that EPS8 and its binding partner IRTKS localized in diffraction limited puncta at future sites of microvilli growth before the F-actin bundling protein ESPIN. EPS8 and IRTKS puncta also remained associated with the distal tips of core actin bundles as these structures elongated. Unexpectedly, we also found that new microvilli commonly emerge from the base of pre-existing protrusions. Finally, we observed that membrane wrapping, membrane-cytoskeleton linking by Ezrin and enrichment of EPS8 and IRTKS are all needed for the survival of nascent microvilli, and that loss of membrane wrapping or tip enriched factors leads to core actin bundle disassembly. These data offer a high resolution molecular and temporal framework for understanding the *de novo* growth of new microvilli.

## RESULTS

### An approach for visualizing the *de novo* growth of individual microvilli

To define the time-course of molecular events that drive microvillar growth, we first established a live imaging approach. Here, we turned to the LLC-PK1-CL4 (CL4) porcine cell culture model, which we recently used to dissect mechanisms of microvillar motility (Meenderink et al., 2019). These cells provide a powerful model for studying apical morphogenesis, as early in differentiation the surface density of microvilli is sparse enough to allow for observations of individual protrusions. CL4 microvilli also extend from the surface at all angles, which enables their visualization in volume projections generated using spinning disk confocal microscopy (Fig. 1A, Video S1, Supp. Fig. 2A). To capture individual growth events, we imaged sub-confluent or recently confluent CL4 cells at 30 second intervals over 30 minutes. Even under these conditions, quantifying individual growth events was limited to a subset of cells and events; e.g. some cells produced few or no *de novo* growth events during the acquisition, overlapping structures precluded measurement, or events did not meet time requirements to gain a full understanding of temporal and molecular dynamics (see Methods). Using mCherry-ESPIN (herein referred to as ESPIN) as a probe for core actin bundles (Loomis et al., 2003), we measured maximum projected ESPIN intensities in the distal half of the growing microvillus. In CL4 cells expressing ESPIN alone, normalized intensity increased, with a growth rate of 0.24 ± 0.12 μm/min and then plateaued after ~5 min (Fig. 1B and C, Supp. Fig 3D). Thus, nascent core actin bundles are assembled on a time-scale of minutes.

**Figure 1.**
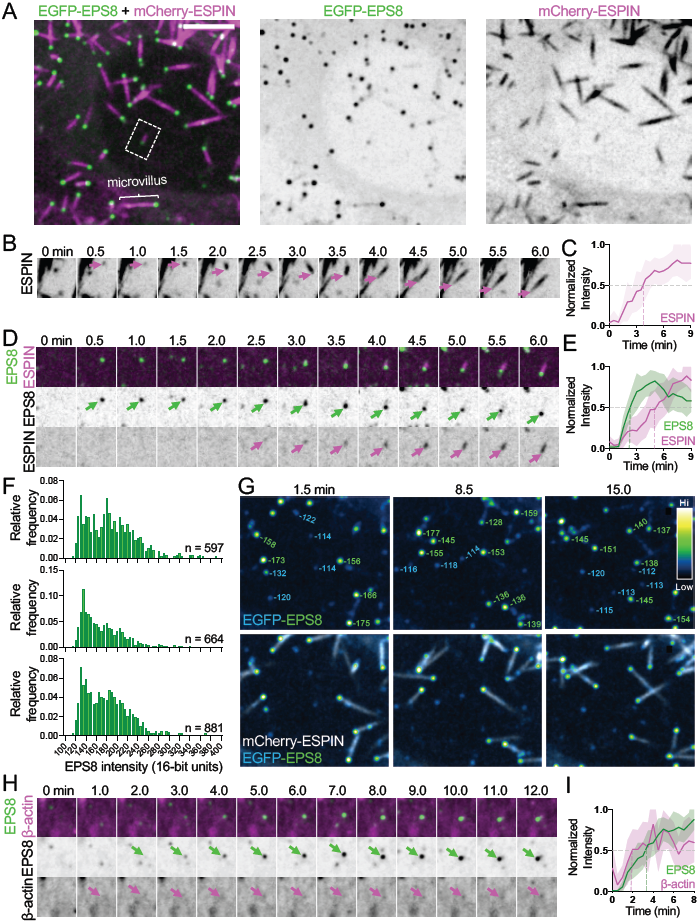
Microvilli emerge from discrete EPS8 puncta on the apical surface of CL4 cells. **(A)** Maximum intensity projection of a CL4 cell displaying microvilli. Core actin bundles are visualized by mCherry-ESPIN (magenta) and EGFP-EPS8 (green). Merge (left), EGFP-EPS8 (middle), mCherry-ESPIN (right). Scale bar = 5 μm. A single microvillus is denoted by the bracket. Dotted box represents microvillus shown in D. **(B)** Montage of a *de novo* microvillus growth event from a CL4 cell overexpressing mCherry-ESPIN alone. Arrows denote the presence of mCherry-ESPIN. Box width = 4 μm. **(C)** Normalized intensity curve of microvillus growth events in CL4 cells expressing mCherry-ESPIN alone, corresponding to montage in D. n = 13 microvilli from 9 cells; t = 0 is defined as −3 frames (1.5 min) before the appearance of the mCherry-ESPIN signal. **(D)** Montage of a *de novo* microvillus growth event from a CL4 cell expressing EGFP-EPS8 and mCherry-ESPIN. Arrows denote the presence of EGFP-EPS8 (green) or mCherry-ESPIN (magenta). Box width = 4 μm. **(E)** Normalized intensity curve of microvillus growth events in CL4 cells expressing EGFP-EPS8 (green) and mCherry-ESPIN (magenta), corresponding to montage in D; n = 14 events from 7 cells. **(F)** EGFP-EPS8 intensity histograms from time-lapse movies of three representative cells. Intensities were calculated from the first frame of each time-lapse; n gives number of EPS8 puncta in each histogram. **(G)** Representative EGFP-EPS8 puncta, pseudocolored by intensity, from the cell shown in A at three different time points. Top row shows EGFP-EPS8 puncta only; the bottom row displays the merge of EGFP-EPS8 and mCherry-ESPIN to differentiate bundle associated and non-bundle associated puncta. Numbers in top row represent raw deconvolved 16-bit intensity units. Green numbers = bundle associated puncta, blue numbers = non-bundle associated puncta. Box width = 14 μm. **(H)** Montage of a *de novo* microvillus growth event in a CL4 cell expressing EGFPEPS8 and mCherry-β-actin. Arrows denote the presence of EGFP-EPS8 (green) or mCherry-ESPIN (magenta). Box width = 4 μm. **(I)** Normalized intensity curve for CL4 cells expressing EGFP-EPS8 (green) and mCherry-β-actin (magenta), corresponding to montage in H. n= 9 growth events from 5 cells. All images shown are maximum intensity projections. Dashed lines for all normalized intensity curves indicate point at which curves cross a normalized intensity of 0.5. Error bars for all curves represent SD. For E and I, t = 0 is defined as −3 frames (1.5 min) before the appearance of the EGFP-EPS8 signal.

### Initiation of microvillus growth coincides with EPS8 enrichment in discrete puncta

Endogenous EPS8 exhibits striking distal tip localization in diverse cell culture models and tissues, in every case examined by us and others (Supp. Figs. 1 and 2)(Croce et al., 2004; Manor et al., 2011a; Postema et al., 2018a; Zwaenepoel et al., 2012). Importantly, endogenous EPS8 localizes to the tips of microvilli in intestinal crypts as well as crypt-like Ls174T-W4 (W4) cells (Supp. Fig. 1A and 2B) suggesting this factor may be involved in early stages of microvilli growth. Moreover, we found endogenous EPS8 distal tip enrichment is reduced in W4 cells when treated with the barbed-end blocking drug, cytochalasin D (Supp. Fig. 2E-G). These data suggest that EPS8 binds directly to filament barbed-ends and therefore, holds the potential to regulate the growth of core actin bundles. With this in mind, we sought to determine if EPS8 arrives before or after the formation of a core actin bundle as marked by ESPIN accumulation. Remarkably, in CL4 cells co-expressing ESPIN and EGFP-EPS8 (herein referred to as EPS8), we found that bright EPS8 puncta appeared minutes before ESPIN accumulation (Fig. 1D and E, Video S2). Microvilli that eventually emerged from these puncta elongated at a rate of 0.11 ± 0.05 μm/min, significantly slower than cells expressing ESPIN alone (Supp. Fig. 3D). Notably, all microvillus growth events observed in our experiments emerged from EPS8 puncta. Analysis of EPS8 puncta intensities revealed distributions that were at least bimodal (Fig. 1F), and core actin bundle associated puncta generally exhibited higher intensities relative to puncta that were not associated with ESPIN positive structures (Fig. 1G, green vs blue numbers). This latter point suggests that puncta likely undergo maturation, accumulating a threshold level of EPS8 before they can support the elongation of microvilli. Although probes that specifically report on the localization of G-actin are not available, co-expression of mCherry-β-actin (labeling G- and F-actin pools) with EPS8 revealed that microvillus growth events were preceded by the coalescence of a diffuse cloud of β-actin, which coincided with the appearance of EPS8 puncta (Fig. 1H and I, Video S3). In combination, these findings lead us to conclude that EPS8 puncta represent “initiation complexes”, which form at the apical surface and incorporate β-actin minutes before core actin bundle assembly begins.

### Mutation of EPS8 actin binding residues impacts initiation complex formation

As specific residues at the C-terminus of EPS8 have been implicated in actin binding (Hertzog et al., 2010), we sought to determine if these motifs are important for EPS8 enrichment in initiation complex puncta or the subsequent growth of nascent microvilli (Fig. 2, Supp. Fig 3). We first asked whether EPS8 localization is dependent on actin binding activity. To address this question, we overexpressed EPS8 mutant constructs in the W4 cell culture model. W4 cells grow brush borders with microvilli that extend parallel to the X-Y image plane, allowing for visualization of fluorescent protein localization along the protrusion axis at ~120 nm resolution when combined with structured illumination microscopy (Grega-Larson et al., 2015; Postema et al., 2018a). Compared to control cells expressing soluble EGFP, cells expressing EPS8 exhibit significant tip enrichment as measured by a distal tip to cytosol ratio (Supp. Fig. 3C). Cells expressing a construct lacking the entire C-terminal actin binding region (EPS8 1-648) or constructs with point mutations in specific actin binding residues (EPS8 V690D L694D, capping defective; LNK758-760AAA or LNK:AAA, bundling defective) (Hertzog et al., 2010), demonstrated impaired tip targeting relative to full length EPS8 (Supp. Fig. 3B and C). We also found that the C-terminal actin binding region (EPS8 849-822) by itself was unable to target to the tips of microvilli (Supp. Fig. 3B and C). These results suggest that, although EPS8 actin binding is necessary for robust tip enrichment, actin binding alone is not sufficient to drive this localization. This point is also consistent with the observation that EPS8 enrichment precedes the bundling of actin filaments by ESPIN, which indicates that EPS8 localization is at least partially independent of actin bundle formation.

**Figure 2.**
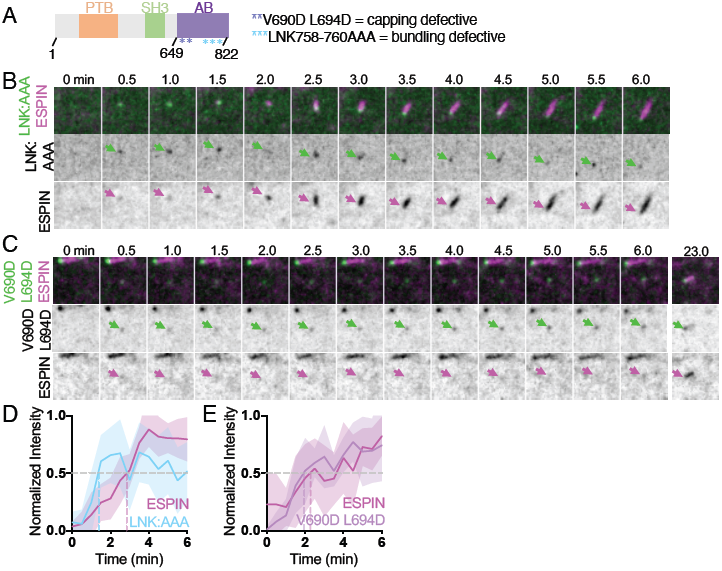
EPS8 actin binding mutants disrupt initiation complex formation and core bundle growth. **(A)** Domain diagram of human EPS8. PTB = phosphotyrosine binding domain, SH3 = src-homology3 domain, AB = actin binding domain. Asterisks represent point mutations in the residues shown. **(B)** Montage of a *de novo* microvillus growth event in cell expressing EGFP-EPS8 LNK758-760AAA (LNK:AAA) and mCherry-ESPIN. Arrows denote the presence of EGFP-EPS8 LNK:AAA (green) or mCherry-ESPIN (magenta). Box width = 4μm. **(C)** Montage of a *de novo* microvillus growth event in cell expressing EGFP-EPS8 V690D L694D and mCherry-ESPIN. Arrows denote the presence of EGFP-EPS8 V690D L694D (green) or mCherry-ESPIN (magenta). Box width = 4 μm. **(D)** Normalized intensity curve of cell expressing EGFP-EPS8 LNK:AAA (light blue) and mCherry-ESPIN (magenta), corresponding to montage in B. n = 11 growth events from 5 cells. **(E)** Normalized intensity curve from cells expressing EGFP-EPS8 V690D L694D (lavender) and mCherry-ESPIN (magenta), corresponding to montage in C. n = 9 growth events from 4 cells. All images shown are maximum intensity projections. For all normalized intensity curves, dashed lines indicate when curves cross a normalized intensity of 0.5. Error bars for all curves shown represent SD. For D and E, t=0 is defined as −3 frames (1.5 min) before the appearance of the EGFP-EPS8 LNK:AAA or EGFP-EPS8 V690D L694D signal.

To determine if mutations in EPS8 actin binding residues impact microvilli growth, we co-expressed the EPS8 LNK:AAA or EPS8 V690D L694D mutants with ESPIN in CL4 cells (Fig. 2). Although appearance of EPS8 LNK:AAA generally preceded ESPIN accumulation, its signal exhibited pronounced intensity fluctuations during the time-course (Fig. 2B and D). Additionally, the microvillus growth rate of cells expressing EPS8 LNK:AAA (0.20 ± 0.15 μm/min) was not significantly different from cells expressing ESPIN alone, suggesting the LNK residues normally serve to limit the speed of microvillus elongation (Supp. Fig. 3D). Moreover, while the appearance of EPS8 V690D L694D coincided with a diffuse cloud of ESPIN, coalescence of this cloud into a linear structure was delayed, in some instances occurring several minutes after the initial appearance of ESPIN signal (Fig. 2C and E,). Cells expressing the EPS8 V690D L694D mutant also exhibited significantly slower microvillus growth rates (0.07 ± 0.32 μm/min) compared to that of cells expressing ESPIN alone (Supp. Fig. 3D). Finally, the EPS8 LNK:AAA mutant exhibited significantly decreased initial mean intensities (measured in the first frame of detectable punctate EGFP signal) compared to EPS8 (Supp. Fig. 3E). Together these fixed and live cell data suggest that the LNK758-760 and V690 L694 actin binding motifs promote accumulation of EPS8 in initiation complexes and efficient formation of core bundles; with the LNK758-760 residues also serving to limit the speed of protrusion growth.

### Nascent microvilli also emerge from pre-existing protrusions

In contrast to *de novo* growth events, we unexpectedly observed that microvilli also grow from pre-existing microvilli. In these cases, a nascent “mother” microvillus gives rise to one or more “daughter” microvilli (Fig. 3A, M = mother, D = daughter). New daughter core bundles grew at angles to the pre-existing mother, giving rise to a structure with a branched or forked appearance. Daughters themselves gave rise to new daughters, which each became separated from the mother bundle (Fig. 3B). Less commonly, we observed daughter microvilli growing from the sides of existing bundles rather than the base (Fig. 3C and Video S4). 3D volume rendering of these structures confirmed that daughters are initially physically linked to a mother core bundle. Intriguingly, the growth of daughter microvilli was marked by small dim puncta of EPS8 that traveled retrograde from the distal tip of an existing mother to a site near its base (Fig. 3A-C). We also noted dim EPS8 puncta localized to the base of microvilli in fixed CL4 cells stained for endogenous EPS8 (Supp. Fig. 2A, top zoom, green arrows). Interestingly, the intensity of EPS8 at the distal tip of a microvillus dropped following the appearance of dim retrograde traveling puncta (Fig. 3D, compare “pre-” vs “post-”). Intensity analysis of distal tip vs. retrograde traveling EPS8 puncta revealed that the EPS8 signal that gives rise to daughter microvilli appears at the expense of EPS8 signal at the distal tip, i.e. EPS8 puncta that give rise to daughter microvilli are initially derived from distal tip puncta (Fig. 3D, Zoom). Based on these data, we propose that formation of daughter microvilli from pre-existing mothers might represent a mechanism for rapidly populating the apical surface with protrusions, one that takes advantage of assembly factors such as EPS8 that are enriched in the vicinity of pre-existing structures.

**Figure 3.**
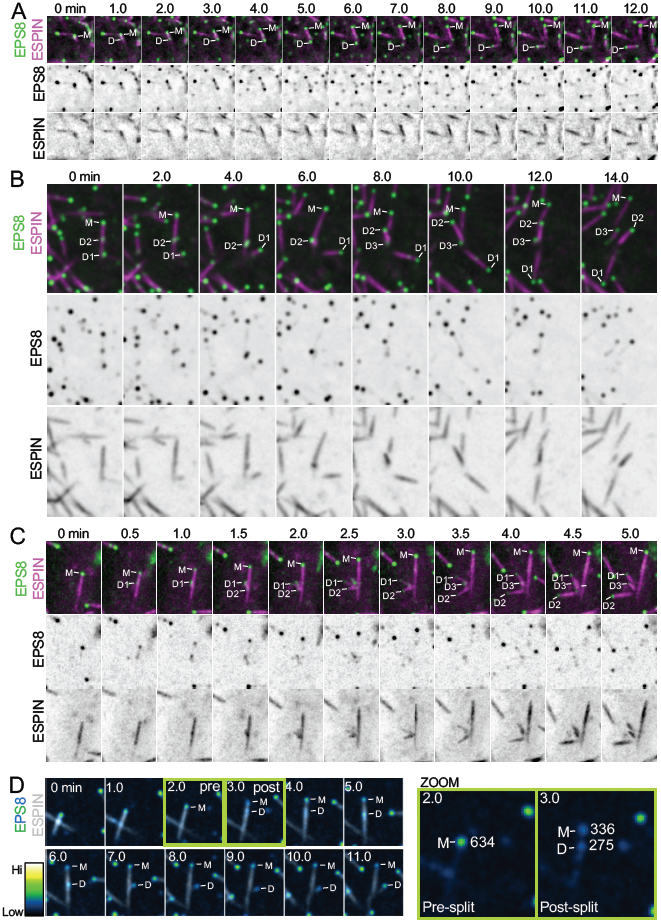
New microvilli also grow from pre-existing microvilli. **(A)** Representative montage of a daughter microvillus growing from the base of an existing mother bundle in a CL4 cell expressing EGFP-EPS8 and mCherry-ESPIN. “M” denotes EPS8 at the tip of the mother microvillus and “D” denotes EPS8 which will eventually mark the tip of a growing daughter microvillus. Box width = 6μm. **(B)** Representative montage of a mother microvillus in a CL4 cell expressing EGFP-EPS8 and mCherry-ESPIN giving rise to multiple daughter microvilli from the the base of the mother bundle. “M” denotes EPS8 at the tip of the mother microvillus and “D1-3” denote EPS8 at the tip of the growing daughter microvilli. Box width = 6.6 um. **(C)** Representative montage of multiple daughter microvilli growing from the side of a mother microvillus in a CL4 cell expressing EGFP-EPS8 and mCherry-ESPIN. “M” denotes EPS8 at the tip of the mother microvillus and “D1-3” denote EPS8 at the tip of the growing daughter microvilli. Box width = 6 μm. **(D)** Montage of a daughter microvillus growth event (left) with EGFP-EPS8 pseudocolored by intensity. The corresponding quantification of EGFP-EPS8 integrated 16-bit intensity pre- and post-splitting of tip localized EPS8 is highlighted (right, Zoom). Note that the total post-split integrated intensity is nearly equal to the pre-split integrated intensity. Green boxes highlight pre- and post-split frames. Box width = 5.6 μm. All images shown are maximum intensity projections.

### IRTKS is also enriched in microvillus growth initiation complexes

Previous work from our group implicated IRTKS as a binding partner of EPS8 and demonstrated that these proteins work together to promote the elongation of microvilli (Postema et al., 2018a). IRTKS possesses both membrane and actin binding motifs through its I-BAR and WH2 domains, respectively, and localizes to the tips of nascent microvilli in the crypts of intestinal organoids and W4 cells in culture (Postema et al., 2018a; Zhao et al., 2011). Based on these properties, we sought to determine if IRTKS, like EPS8, accumulates in the initiation complexes that give rise to microvilli. EGFP-IRTKS (herein referred to as IRTKS) exhibited enrichment in discrete puncta at the tips of existing microvilli, and also demonstrated some lateral localization as well, as shown previously (Fig. 4A) (Meenderink et al., 2019). Like EPS8, IRTKS localized to sites of microvilli growth minutes before core bundle formation as marked by ESPIN accumulation (Fig. 4B and D a Video S5). The growth of daughter microvilli was also marked by small dim IRTKS puncta that were laterally associated with pre-existing mother core bundles (Fig. 4C). These results demonstrate that the EPS8 binding partner IRTKS also accumulates in initiation complexes at the earliest stages of microvillus growth.

**Figure 4.**
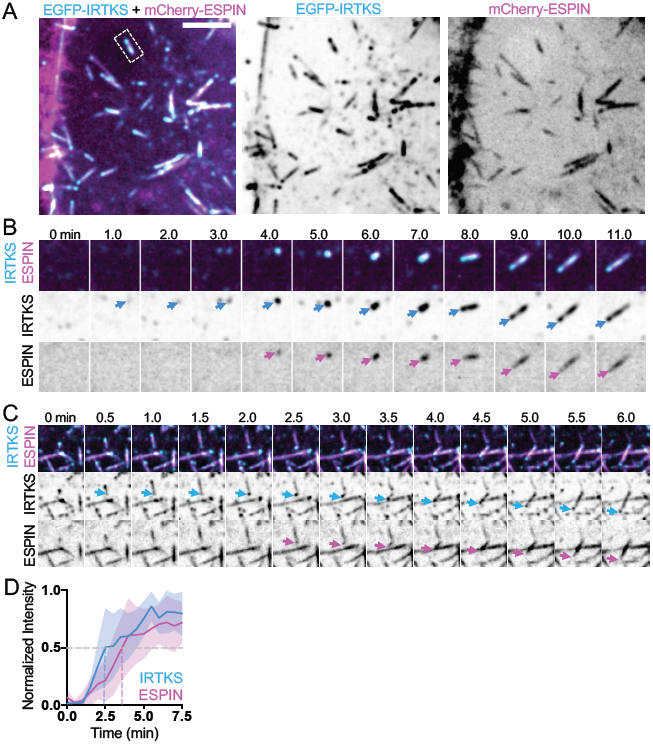
IRTKS is also enriched in microvillus initiation complexes. **(A)** Maximum intensity projection of CL4 cell expressing mCherry-Espin (magenta) and EGFP-IRTKS (cyan). Merge (left), EGFP-IRTKS (middle), mCherry-ESPIN (right). Dashed box indicates microvillus shown in B. Scale bar = 5 μm. **(B)** Montage of a *de novo* microvillus growth event in a cell expressing EGFP-IRTKS and mCherry-ESPIN. Arrows denote the presence of EGFP-IRTKS (cyan) or mCherry-ESPIN (magenta). Box width = 4μm. **(C)** Montage of daughter microvillus growing from pre-existing mother bundle in a cell expressing EGFP-IRTKS and mCherry-ESPIN. Arrows denote EGFP-IRTKS (cyan) and mCherry-ESPIN (magenta) of the daughter microvillus. Box width = 6 μm. **(D)** Normalized intensity curve of CL4 cells expressing EGFP-IRTKS and mCherry-ESPIN; n = 9 events from 4 cells. t = 0 is defined as −3 frames (1.5 min) before the appearance of the EGFP-IRTKS signal. Dashed lines indicate point at which curves cross a normalized intensity of 0.5. Error bars for all curves show SD. All images shown are maximum intensity projections.

### Core actin bundle elongation coincides with plasma membrane wrapping

As microvilli are plasma membrane wrapped protrusions, and because IRTKS directly interacts with the membrane through its I-BAR domain (Saarikangas et al., 2009), we sought to determine the timing of membrane wrapping relative to the formation of initiation complexes and the subsequent growth of the core bundle. To test whether enrichment of IRTKS precedes membrane encapsulation, we incubated cells co-expressing IRTKS and ESPIN with the live cell plasma membrane label CellMask-DeepRed (herein referred to as CellMask) (Fig. 5A and D, Video S6). Individual microvilli typically exhibited CellMask enrichment at the distal end, which we interpreted as membrane encapsulation (Fig. 5F). During *de novo* growth events, IRTKS enriched in initiation complex puncta prior to the accumulation of ESPIN and CellMask, which both increased in parallel (Fig. 5A and D). Additionally, we found that the daughter microvilli that grow from pre-existing mothers also become wrapped in membrane (Fig. 5B). Because previous studies established the importance of membrane-cytoskeleton linker proteins (i.e. factors that simultaneously bind membrane and F-actin) in normal microvilli morphology (Pelaseyed and Bretscher, 2018) we next sought to determine if the membrane wrapping during microvilli growth was driven by the accumulation of such a factor. Here, we focused on the Ezrin-Radixin-Moesin (ERM) family protein EZRIN, a well-established membrane-cytoskeleton linker that is known to mechanically stabilize epithelial microvilli (Pelaseyed and Bretscher, 2018). Live imaging revealed that EZRIN-EGFP (herein referred to as EZRIN) and ESPIN accumulated in parallel during microvillus growth (Fig. 5C and E and Video S7). These data suggest that core bundle formation and membrane wrapping driven by ERM proteins are parallel events in microvillus growth, which follow the appearance of initiation complexes at the apical surface.

**Figure 5.**
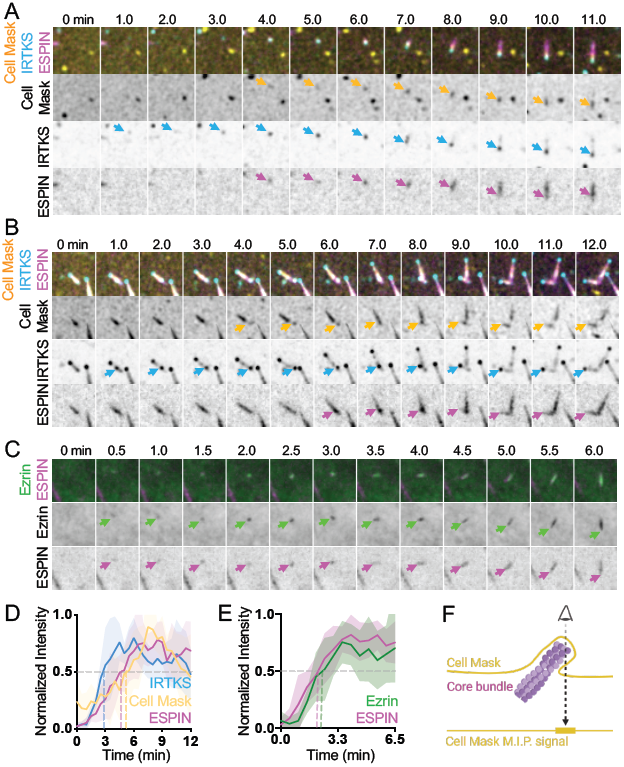
Membrane encapsulation occurs in parallel with core bundle formation. **(A)** Montage of *de novo* microvillus growth event in a CL4 cell expressing EGFP-IRTKS and mCherry-ESPIN and stained with CellMask-DeepRed to mark the plasma membrane. Arrows denote the onset of membrane wrapping visualized by CellMask-DeepRed (yellow), or the presence of EGFP-IRTKS (cyan) and mCherry-ESPIN (magenta). Box width = 4μm. **(B)** Representative montage of daughter microvillus formation in a cell expressing EGFP-IRTKS and mCherry-ESPIN and stained with CellMask-DeepRed. Arrows denote the onset of membrane wrapping as visualized by CellMask-DeepRed (yellow), or the presence of EGFP-IRTKS (cyan) and mCherry-ESPIN (magenta) of the daughter microvillus. **(C)** Montage of a *de novo* microvillus growth event in a CL4 cell expressing Ezrin-GFP and mCherry-ESPIN. Arrows denote the presence of Ezrin-GFP (green) or mCherry-ESPIN (magenta). Box width = 4μm. **(D)** Normalized intensity curves of cells expressing EGFP-IRTKS and mCherry-ESPIN and stained with CellMask-DeepRed corresponding to montage in A. n = 6 growth events from 3 cells. **(E)** Normalized intensity curves of cells expressing Ezrin-GFP and mCherry-ESPIN corresponding to montage in C. n = 11 growth events from 8 cells. **(F)** Schematic representation of a microvillus stained with cell mask and the resulting maximum intensity projected signal. All images shown are maximum intensity projections. Dashed lines for normalized intensity curves indicate point at which curves cross a normalized intensity of 0.5. Error bars for all curves show SD. For D and E, t = 0 is defined as −3 frames (1.5 min) before the appearance of the EGFP-IRTKS or Ezrin-GFP signal.

### Core bundles cannot survive without plasma membrane wrapping

Analysis of our time-lapse data revealed that a subset of nascent microvilli “collapsed”, which was characterized by loss of the core actin bundle and associated factors. Analysis of cells expressing EPS8 and ESPIN revealed that collapse events are typically preceded by loss of EPS8 puncta from the distal tips (Fig. 6A and C and Video S8), which was immediately followed by a decrease in core bundle length (Fig. 6A and C). We also observed instances where core bundles appeared to “break”; in these cases, we noted that the core bundle remnant from the original microvillus persisted and maintained distal tip enriched EPS8 signal. However, the newer core bundle fragment lacked EPS8 tip enrichment and quickly collapsed (Supp. Fig. 5A). We observed the same general pattern with cells co-expressing IRTKS and ESPIN, whereby loss of punctate IRTKS signal was followed by a decrease in microvilli length (Fig. 6B and D). These results suggest that loss of EPS8 and IRTKS from the distal tip may destabilize nascent microvilli.

**Figure 6.**
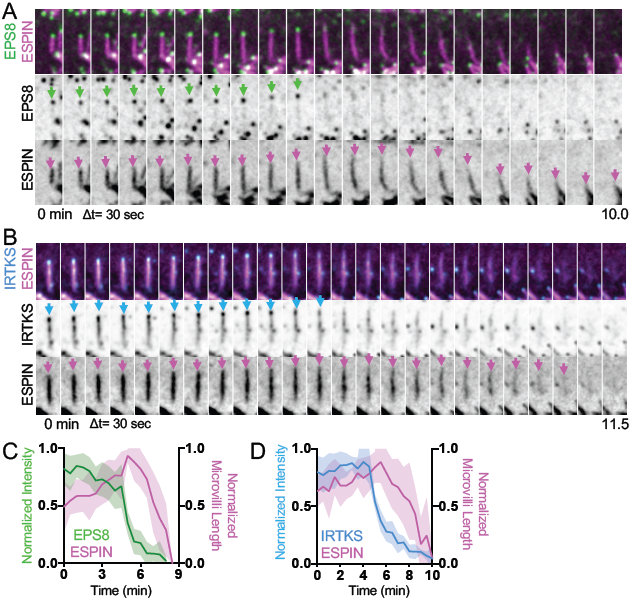
Microvilli cannot survive without EPS8 and IRTKS at the tips. **(A)** Montage of a microvillus collapse event on the surface of a CL4 cell expressing EGFP-EPS8 and mCherry-ESPIN. Green arrows denote tip targeted EGFP-EPS8 signal. Magenta arrows denote the core actin bundle. Box width = 2.5μm. **(B)** Montage of microvillus collapse event in a cell expressing EGFP-IRTKS and mCherry-ESPIN. Cyan arrows denote tip targeted EGFP-IRTKS signal. Magenta arrows denote the core actin bundle. Box width = 2.5 μm. **(C)** Quantification of EGFPEPS8 intensity and microvilli length as represented by montage in A. n = 10 events from 6 cells. **(D)** Quantification of EGFP-IRTKS intensity and microvilli length as represented in B. n = 5 events from 2 cells. For C and D, t = 0 is defined as −10 frames (5 min) before the drop in EGFPEPS8 or EGFP-IRTKS intensity at the tips of microvilli. All images are represented as maximum intensity projections. Error bars for all curves show SD.

These observations prompted us to investigate upstream events that could potentially lead to loss of tip targeted factors and subsequent microvilli collapse. As IRTKS and EPS8 are enriched at the interface of the apical membrane and actin core bundle, we again used CellMask to visualize the plasma membrane during collapse events. Interestingly, we found that CellMask signal, which we interpret as protruding membrane, is lost prior to collapse (Fig. 7A-D). Loss of CellMask signal was followed by loss of punctate IRTKS signal and a subsequent decrease in microvillus length as marked by ESPIN (Fig. 7A and C, Supp. Fig 4B and Video S9). As the plasma membrane is linked to the underlying actin cytoskeleton through membrane-cytoskeleton linkers, we sought to determine if loss of membrane encapsulation was linked to a loss of EZRIN signal from nascent microvilli. Indeed, we observed that CellMask and EZRIN signals decreased in parallel, before a measurable decrease in microvilli length as marked by ESPIN (Fig. 7B and D, Supp. Fig 4C and Video S10). These results suggest that microvilli cannot survive without membrane wrapping, loss of which is paralleled by loss of membrane-cytoskeleton linking, followed by loss of distal tip factors, and ultimately collapse of the core bundle. These findings also reveal that early in epithelial differentiation, nascent microvilli undergo cycles of growth and collapse, which likely offer a degree of plasticity during the complex process of apical morphogenesis.

**Figure 7.**
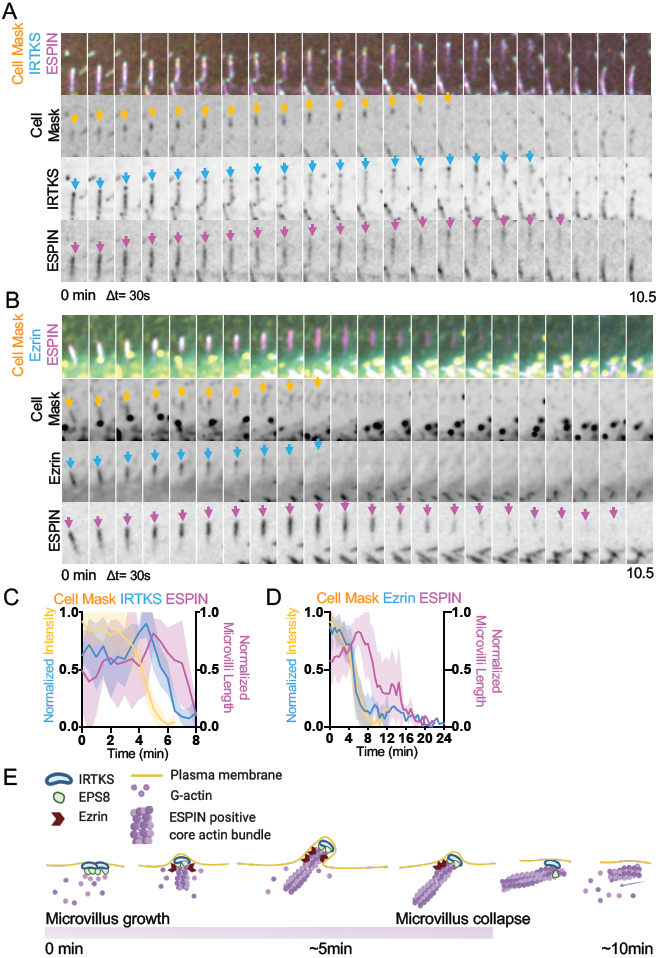
Membrane encapsulation is essential for microvillus survival. **(A)** Montage of a microvillus collapse event in a cell expressing EGFP-IRTKS and mCherry-Espin, stained with CellMask-DeepRed. Z-coded ESPIN is included to show the core bundle does not drift out of plane. Box width = 2.5 μm. **(B)** Montage of a microvillus collapse event in a cell expressing Ezrin-GFP and mCherry-Espin, stained with CellMask-DeepRed. Z-coded ESPIN is included to show the core bundle does not drift out of plane. Box width = 2.5 μm. Quantification of EGFP-IRTKS and CellMask-DeepRed intensity and microvilli length as represented in A. n = 4 events from 4 cells. **(D)** Quantification of Ezrin-GFP and CellMask-DeepRed intensity and microvilli length as represented in B. This plot contains averaged traces of varying lengths that range in duration from 6.5 - 24 minutes (EZRIN), 6 - 12 min (Cell Mask) and 7.5 - 22 min (ESPIN) with n = 15 events 10 cells. Error bars in all curves represent SD. For C and D, t=0 is defined as −10 frames (5 min) before the drop in EGFP-IRTKS or Ezrin-GFP intensity at the tips of microvilli. All images are represented as maximum intensity projections. Error bars for all curves represent SD. **(E)** Model of *de novo* microvillus growth and collapse. Formation of initiation complexes composed of EPS8 and IRTKS coincide with local enrichment of G-actin. Formation of these initiation complexes is followed by assembly of ESPIN positive core actin bundles, recruitment of Ezrin, and plasma membrane protrusion. The process of *de novo* growth occurs on the scale of ~5 minutes. During microvillus collapse, nascent bundles lose membrane wrapping and enrichment of EZRIN in parallel. This loss of membrane wrapping destabilizes initiation complex proteins EPS8 and IRTKS, which leads to collapse of the ESPIN positive core actin bundle. The process of collapse also occurs on the scale of ~5 minutes. This model cartoon was created using BioRender.com.

## DISCUSSION

Here we employed a live imaging approach to define a time-course for the recruitment of factors that drive the growth and stabilization of new microvilli. Using CL4 cells as an epithelial model system, our experimental goal was to capture the earliest events underlying biogenesis of these protrusions. The resulting spatially and temporally resolved datasets allow us to stage the appearance of a core set of microvillar components including G-actin, tip enriched actin regulatory factors, filament bundling proteins, membrane-cytoskeleton linkers and the plasma membrane during this dynamic process (Fig. 7E). The growth of a new microvillus takes place over the course of several minutes, similar to the timeframe reported for microvilli in Xenopus kidney A6 cells (Gorelik et al., 2003). The emergence of a nascent protrusion is first signaled by the enrichment of distal tip factors EPS8 and IRTKS, in diffraction limited puncta at the apical surface. EPS8 enrichment is also accompanied by a moderate accumulation of G-actin. Because the appearance of EPS8/IRTKS puncta preceded the formation of microvilli observed in our assays, we propose that these factors are components of the electron dense tip complex, which has long been proposed to function in initiating protrusion growth (Tilney and Cardell, 1970). The tip enrichment of EPS8 is at least partially driven by binding to actin, as mutations in actin binding residues interfered with initiation complex formation, and EPS8 puncta not associated with actin bundles typically exhibited lower intensities. Furthermore, mutation of the V690 and L694 residues resulted in delayed core bundle formation suggesting direct involvement of EPS8 in the growth of these structures. Several minutes after the appearance of EPS8/IRTKS containing initiation complexes, core actin bundles form as indicated by accumulation of the parallel bundling protein ESPIN. Parallel enrichment of ESPIN with the CellMask membrane probe and the membrane-cytoskeleton linking protein Ezrin suggests that actin core bundles are assembled in close proximity to the apical surface, where they are quickly or immediately encapsulated in plasma membrane. The time course of molecular recruitment described here provides a framework for understanding how cells control the growth of surface features, and may also apply to related finger-like protrusions including filopodia and stereocilia.

In addition to the *de novo* microvilli growth events described above, we also observed the formation of daughter microvilli from a pre-existing mother protrusions. Formation of a new daughter core bundle was preceded by enrichment of EPS8 and IRTKS at the site of lateral growth from the mother. Relative to *de novo* assembly, such mother/daughter growth might be energetically favorable as materials for growing a new microvillus, namely initiation complex proteins and actin subunits, are enriched near pre-existing protrusions. Moreover, the physical obstacle presented by plasma membrane bending could be partially alleviated by the mother protrusion. We propose that in combination with *de novo* growth, mother/daughter growth allows epithelial cells to quickly populate their apical surface with microvilli. This would be particularly advantageous in the intestinal epithelium, where cells rapidly increase the number of apical microvilli as they migrate from stem cell containing crypts out to the villus surface (Specian and Neutra, 1981; van Dongen et al., 1976). Although daughter microvilli eventually separate from mother protrusions, the mother/daughter growth mechanism initially gives rise to structures with a forked appearance. Such structures are strikingly reminiscent of the “forked microvilli” phenotype captured using TEM many years ago (Tilney and Cardell, 1970). In that study, hydrostatic pressure was used to efface microvilli from the apical surface of small intestine explants. Microvilli were then allowed to re-assemble before fixation, resulting in some structures with a split or “forked” appearance. Remarkably, the authors noted electron dense plaques on the lateral surface of the new core bundles, which were hypothesized to represent precursors of forked microvilli. One intriguing possibility is that the lateral plaques observed by Tilney and Cardell represent protein complexes containing EPS8 and IRTKS, which would be consistent with what we visualized during mother/daughter growth events in our live time-lapse data. Moreover, as the Tilney and Cardell study used intestinal epithelial samples from salamander and the observations we report here focused on cells derived from porcine kidney, we suspect that the mother/daughter mechanism of microvilli growth is conserved across species and tissue types.

Interestingly not all nascent microvilli survive and a notable fraction of the protrusions observed in our time-lapse datasets collapsed during the period of observation (Fig. 7E). Collapse of a microvillus is predicted by loss of plasma membrane wrapping, which occurs in parallel with decreased membrane-cytoskeleton linking by EZRIN. Loss of membrane encapsulation is followed by loss of tip targeted EPS8 and IRTKS, which in turn is ultimately followed by collapse of the core actin bundle. Thus, in order to survive, core bundles must remain intimately linked to and partially encapsulated by the plasma membrane. What leads to the loss of membrane-cytoskeleton linking and plasma membrane wrapping is not clear. As our time-lapse observations take place at the earliest stages of apical surface maturation, one possibility is that the cells are not yet expressing the factors needed to stabilize microvilli at high levels. This would be consistent with previous studies showing that a number of microvillus resident proteins gradually accumulate throughout the time course of differentiation (Crawley et al., 2014a; Heintzelman and Mooseker, 1990). Alternatively, collapse events in a particular region might indicate that local membrane composition or mechanics are not favorable for protrusion. Yet another possibility is that microvilli are by default destined for collapse and only a subset are selected for stabilization. For example, adherence with neighboring protrusions driven by tip-targeted protocadherins (CDHR5 and CDHR2) could provide a mechanical capture mechanism that permits the long-term survival of new microvilli as we recently proposed (Meenderink et al., 2019). Such dynamics might also allow for error correction, where new microvilli that arise in the wrong location, i.e. far from neighboring and potentially adherent protrusions, are eliminated through collapse. Molecular subunits made available during these events could then be used to regrow a protrusion in a more crowded location, in turn accelerating brush border maturation. Events where microvilli collapse precedes the formation of multiple new daughter microvilli from remnants of EPS8 puncta are consistent with the general idea of building block recycling in this system (Supp. Fig. 5B).

While this study offers a temporally resolved molecular framework for understanding the growth of new microvilli, important questions remain unanswered. One question of longstanding interest pertains to mechanisms that control the number of actin filaments per core bundle. Microvilli in mature transporting epithelia exhibit a highly stereotyped number of actin filaments (~30-40) per core bundle (Ohta et al., 2012), suggesting that cells enlist mechanisms to control the stoichiometry of bundle components. One attractive hypothesis is that proteins in the distal tip complex control filament numbers (Tilney and Cardell, 1970). If EPS8 does interact directly with core bundle actin filament barbed-ends, the number of EPS8 molecules per tip puncta might ultimately dictate the number of actin filaments per core bundle. Interestingly, core bundle associated EPS8 puncta are stereotyped in intensity/size, supporting the general idea that EPS8 stoichiometry in these complexes is regulated. Although the EPS8 overexpression approach used here precludes absolute molecular counting measurements, future studies taking advantage of endogenously tagged models should allow for quantitative dissection of distal tip puncta composition and stoichiometry.

Another important unanswered question relates to how distal tip enriched factors may be recruited to the plasma membrane. Epithelial cells are characterized by polarized separation of phosphatidylinositols, with phosphatidylinositol 4,5 bisphosphate (PI(4,5)P_2_) enriched at the apical surface and phosphatidyl inositol 3,4,5-trisphosphate (PI(3,4,5)P_3_) enriched basolaterally (Gassama-Diagne et al., 2006; Martin-Belmonte et al., 2007). Although the emergence of EPS8 and IRTKS puncta appears to be stochastic in nature, the local enrichment of signaling lipids such as PI(4,5)P_2_, could be involved in recruitment of these factors. Both EPS8 and IRTKS contain structural motifs that make this a reasonable possibility. For instance, the EPS8 PTB domain shares sequence similarity with the PTB domain of Dab1, which has been shown to bind PI(4,5)P_2_ (Uhlik et al., 2005). Similarly, other proteins of the I-BAR family such as MIM and IRSP53, have also been shown to directly bind PI(4,5)P_2_ (Mattila et al., 2007; Saarikangas et al., 2009). Future time-resolved studies must focus on defining the nature of EPS8 interactions with membrane lipids, and whether apical localization of EPS8 and IRTKS is controlled by upstream enrichment of specific lipid species.

In summary, this study is the first to capture the molecular details of microvilli biogenesis using a live imaging approach. The resulting time-lapse datasets allow us to construct a temporal framework for key factors involved in microvillus growth, and offer insight into the nature of apical surface dynamics at early timepoints in epithelial differentiation. These results also hold implications for understanding the assembly and dynamics of related actin supported protrusions. For example, classic ultrastructural studies of early stage chick embryos revealed that the formation of stereocilia is preceded by the growth of microvilli (Tilney et al., 1992). A fraction of these microvilli are resorbed, and the remaining protrusions are believed to mature into stereocilia arranged in the characteristic staircase pattern. The collapse dynamics we report here may offer a mechanism to explain microvillar resorption during sensory epithelium differentiation.

## ACKNOWLEDGEMENTS

The authors would like to thank all members of the Tyska laboratory, members of the Vanderbilt Microtubules and Motors Club, the Epithelial Biology Center, and the Cellular, Biochemical, and Molecular Sciences Training program for feedback and guidance. Microscopy was performed in part by the Vanderbilt Cell Imaging Shared Resource. This work was supported by the Vanderbilt Cellular, Biochemical and Molecular Sciences Training Grant 5T32GM008554-25 (I.M.G.), the NIH NIDDK National Research Service Award F31DK122692 (I.M.G.), American Heart Association Predoctoral Fellowship (M.M.P.), Program in Developmental Biology Training Grant 5T32HD007502-20 (CSC), NIH grant T32-A1007474 (L.M.M.), Department of Veterans Affairs Career Development Award 1IK2-BX004885 (L.M.M.), and NIH grants R01-DK111949 and R01-DK095811 (M.J.T.).

## AUTHOR CONTRIBUTIONS

I.M.G. and M.J.T. designed the experiments. I.M.G. performed all experiments, analyzed the data, and prepared the figures. L.M.M. provided expertise with experimental methodology and significant conceptual insight. M.M.P. and C.S.C. assisted with data collection. I.M.G. and M.J.T. wrote the manuscript. All authors contributed to revising the manuscript.

## DECLARATION OF INTERESTS

The authors declare no competing interests.

## MATERIALS AND METHODS

### Cell culture

LLC-PK1-CL4 (porcine kidney proximal tubule) and Opossum Kidney (OK) cells were cultured in DMEM with high glucose and 2mM L-glutamine supplemented with 10% fetal bovine serum (FBS) and 1% glutamine. Caco_2BBE_ cells were cultured in DMEM with high glucose and 2mM L-glutamine supplemented with 20% FBS and 1% glutamine. Ls174T-W4 cells (female *Hs* colon epithelial cells) were cultured in DMEM with high glucose and 2 mM L-glutamine supplemented with 10% tetracycline-free FBS, 1% glutamine, G-418 (1 mg/ml), blasticidin (10 μg/ml), and phleomycin (20 μg/ml). This cell line was obtained directly from Dr. Hans Clevers (Utrecht University, Netherlands) and has not been additionally authenticated beyond confirmation of polarization with the addition of doxycycline. All cultures were maintained at 37°C with 5% CO_2_.

### Cloning and constructs

pLVX-mCherry-ESPIN, EGFP-EPS8, and EGFP-IRTKS were described previously (Meenderink et al., 2019; Postema et al., 2018a), whereas Ezrin-EGFP was a gift from Dr. Stephen Shaw (NIH). The full length EGFP-EPS8 construct used for most experiments described herein contained two amino acid substitutions in the C-terminus (E818D and S820C) and made use of the stop codon in the plasmid backbone ~15 amino acids downstream of the native stop. All other EPS8 constructs are based on the sequence accession AAH30010.1 except for the indicated mutations or truncations. EGFP-EPS8 1-648 (primers 5’→3’ forward atgaatggtcatatttctaatcatccc; 5’→3’ reverse ttatgacacaggaacaggtgctgg) and EGFP-EPS8 649-822 (primers 5’→3’ forward aaggtcccagcaaatataacacgtca; 5’→3’ reverse ttagtgactgcttccttcatcaaaagattcc) constructs were generated by PCR, topo-cloned into the pCR8 entry vector (Invitrogen), and then shuttled into a gateway adapted EGFP-C1-GW vector via the LR recombination reaction. Point mutagenesis was performed on a pCR8-EPS8 template, and then shuttled into EGFP-C1-GW vector to create EGFP-EPS8 V690D L694D (primers 5’→3’ forward gaaatctcagatggaggaagaccaagatgaagacatccacagactgaccattg; 5’→3’ reverse caatggtcagtctgtggatgtcttcatcttggtcttcctccatctgagatttc) and EGFP-EPS8 LNK758-760AAA (primers 5’→3’ forward ggtgcacaacttttctctgccgctgcggatgaactgaggacagtctgccc; 5’→3’ reverse gggcagactgtcctcagttcatccgcagcggcagagaaaagttgtgcacc). To create pLVX-mCherry-βactin, the empty pLVX-puro vector was first created by digesting pLVX-AcGFP-C1 (Takara) with XhoI and NdeI, and then ligating a region of the CMV promoter cut during digestion. This region was amplified using primers (5’→3’ forward cccccgcccattgacgtcaataatgacg and 5’→3’ reverse taagcactcgagggtggcgaccggtagat) and digested with XhoI and NdeI. mCherry-βactin was generated by PCR using primers (5’→3’ forward taagcagaattcgccaccatggtgagcaagg; 5’→3’ reverse taagcagggcccctagaagcatttgcggtggacga) and ligated into the pLVX-puro backbone after digestion of the backbone and insert with EcoRI and ApaI. Constructs created for this study were confirmed via sequencing.

### Stable cell line generation and transient transfection

#### Stable cell line generation

To create cells stably expressing two fluorescent proteins, LLC-PK1-CL4 were first transiently transfected with EGFP tagged constructs and subsequently selected with 1mg/mL G-418 for approximately two to four weeks (EGFP-EPS8, EGFP-IRTKS, EGFPEPS8LNKAAA, EGFP-EPS8 V690D L694D). Cells were then transduced with lentivirus expressing mCherry-ESPIN or mCherry-βactin to make the double stable cell line. Lentivirus was generated using either HEK293T or HEK293FT cells co-transfected with pLVX-mCherry-ESPIN or pLVX-mCherry- βactin, and psPAX2 and pMD2.G using lipofectamine 2000, according to the manufacturer’s instructions. Media was changed 16 hours after transfection, and virus was collected 72 and 96 hours after transfection and concentrated using Lenti-X concentrator (Clontech). Cells were then transduced with virus after growing to 75% confluency and supplemented with 10μg/mL polybrene. Media was changed one day after transduction and cells were placed under selection with puromycin (10μg/mL concentration) 72 hours after transduction. Established double stable cell lines were maintained in 1mg/mL G-418 and 10μg/mL concentration puromycin. Cells expressing mCherry-ESPIN were used previously (Meenderink et al., 2019) and were created by transient transfection with pmCherry-ESPIN selected with 1 mg/mL G-418, sorted, and maintained in selection with 1 mg/mL G-418.

#### Transient transfection

W4 cells were seeded in 6 well plates and transfected at ~60% confluency, using Lipofectamine 2000 according to manufacturer’s instructions. The following day, cells were re-plated on acid-washed glass coverslips and induced to form brush borders using 1μg/ml doxycycline for 16hr before fixation.

### Immunofluorescence

#### Cell Culture

Cells were rinsed 1x in prewarmed PBS, and fixed by incubating in prewarmed 4% paraformaldehyde for 15 min at 37°C. Cells were then rinsed 3x for 5 minutes with PBS and permeabilized in 0.1% TritonX-100 for 15 min at room temperature. Cells were then rinsed with PBS 3x for 5 minutes and then blocked in 5% BSA for 1hr at 37°C. Following blocking, cells were briefly rinsed 1x with PBS, and incubated with 1° antibodies for 1hr at 37°C (mouse anti-EPS8, BD Transduction Laboratories Cat#610144 1:400; Chicken IgY anti-GFP, Cat#GFP-1020 Aves Labs 1:200). Cells were then rinsed 4x for 5 minutes with PBS and incubated with 2° antibodies (Goat anti-mouse Alexa Fluor 488 F(ab’)2 fragment, Invitrogen Ref A11017, 1:1000 dilution; Goat anti-mouse Alexa Fluor 568 F(ab’)2 fragment Ref A11019, 1:1000 or Goat-anti chicken Alexa Fluor 488 IgG (H+L), Invitrogen Ref A11039, 1:1000 dilution) and Alexa-Fluor 568 Phalloidin (Invitrogen Ref A12380) or Alexa-Fluor 488 Phalloidin (Invitrogen Ref A12379) at 1:200 dilution for 30 minutes at room temperature. Following 2° antibody incubation, cells were rinsed 4x for 5 minutes in PBS and mounted on glass slides using ProLong Gold antifade reagent (Invitrogen).

#### Tissue

Paraffin-embedded tissue sections of mouse WT small intestinal swiss rolls and kidney sections were deparaffinized using Histo-Clear solution (Fisher) and rehydrated in a descending graded ethanol series (100%, 95%, 90%, 70%, 50%) and rinsed with PBS 3x for 5 min. Slides were then subject to an antigen retrieval step consisting of boiling for 1 h in a solution of 10 mM Tris (pH 9.0) and 0.5 mM ethylene glycol-bis(β-aminoethyl ether)-*N,N,N',N'*-tetraacetic acid (EGTA). Slides were then washed in PBS 3x and blocked with 5% normal goat serum (NGS) overnight at 4°C. After blocking, slides were briefly rinsed with PBS and stained overnight at 4°C with antibodies (mouse anti-EPS8, BD Transduction Laboratories Cat#610144, 1:200; rabbit anti-Villin 1:50, Santa Cruz Cat# SC28283) in 1% NGS in PBS. After being washed with PBS four times, samples were incubated with secondary antibodies (Goat anti-mouse Alexa Fluor 488 F(ab’)2 fragment, 1:1000; anti-Rabbit H+L Alexa Fluor 568 1:1000) in 1% NGS in PBS for 2h at RT. Slides were then washed four times with PBS, dehydrated in an ascending ethanol series (50%, 70%, 90%, 95%, 100%), and mounted using ProLong Gold Antifade reagent (Life Technologies).

### Cytochalasin D treatment

Wild-type W4 cells were treated with 1μg/ml doxycycline for 16hr prior to cytochalasin D treatment to induce brush border formation. Cells were then treated with 500nM cytochalasin D or an equivalent volume of DMSO diluted in W4 media for 30 minutes. Cells were then fixed in 4% paraformaldehyde and stored at 4°C in 1x PBS prior to staining.

### Live cell membrane labeling

CL4 cells were incubated with a 1.5x concentration of CellMask-DeepRed plasma membrane dye (Molecular Probes) in Fluorobrite Medium (Gibco) supplemented with 10% FBS and 1% L-glutamine for 10 min at 37°C. Cells were subsequently imaged in supplemented Fluoroborite Medium to minimize background fluorescence.

### Microscopy

#### Microscopes

Laser scanning confocal microscopy was performed using a Nikon A1R or A1 laser-scanning confocal microscope equipped with 488 and 561 nm excitation LASERs, and 100x/1.49 NA TIRF, 40x/1.3 NA oil, and 25x/1.05 NA silicon immersion objectives. Structured Illumination Microscopy was performed using an N-SIM (Nikon Instruments) equipped with Andor DU-897 EMCCD camera with four color excitation lasers (405 nm, 488 nm, 561 nm, and 647 nm), and a 100x/1.49 NA TIRF objective. All images were reconstructed using Nikon Elements software with matching reconstruction parameters. Spinning Disk Confocal Microscopy was performed using a Nikon Ti2 Nikon Ti2 inverted light microscope equipped with a Yokogawa CSU-X1 spinning disk head, Andor DU-897 EMCCD camera or a Photometrics Prime 95B sCMOS camera, 488 nm and 561 nm excitation lasers, and a 100x/1.49 NA TIRF objective. Cells were imaged in a stage top incubator maintained at 37°C with 5% CO2 (Tokai Hit). *Live cell imaging routine*: In order to capture microvilli growth and collapse events, subconfluent to recently confluent (~confluent the day of imaging) CL4 cells plated on plasma cleaned 35-mm glass bottom dishes (MatTek) were imaged at 30 second intervals for 30 minutes in CL4 media. CL4 cells expressing EGFP-EPS8 and mCherry-βactin were plated on plasma cleaned 35-mm glass bottom dishes coated with 50 μg/mL laminin to minimize cell lifting during the acquisition. Z stacks of ~3-5μm were collected depending on cell thickness and sampled using a 0.18μm step size using a Triggered NIDAQ Piezo Z driver. Image acquisition was performed using Nikon Elements software. Movies with intensities compared across conditions were collected using matching laser power and exposure times.

### Quantification of fluorescence microscopy images

Measurement of microvilli growth events was performed manually in FIJI. All measurements were performed on z-projected images deconvolved using the Richardson-Lucy algorithm with 20 iterations in Nikon Elements Software. For EGFP and mCherry co-expressing cells, t=0 was defined as −3 frames before the appearance of the EGFP signal (EGFP-EPS8, EGFPEPS8LNK:AAA, EGFP-EPS8 V690D L694D, EGFP-IRTKS, Ezrin-GFP). The same ROI was then used to track the mCherry (ESPIN or βactin) signal of the distal to middle region of the growing microvillus. ROIs of approximately the same size were used between independent growth events. For cells expressing mCherry-ESPIN only, t=0 is defined as −3 frames before the appearance of the mCherry signal, with a circular ROI used to measure the distal to middle region of the growing microvillus. For *de novo* growth, events were excluded if there were less than 3 frames before the appearance of the EGFP signal, if the microvillus collapsed less than 6 min after formation, if a single bundle immediately gave rise to multiple microvilli, or if significant overlap with adjacent microvilli confounded analysis. Using Prism V.7 (GraphPad), mean intensity values were first normalized to account for intensity differences between microvilli, with the smallest value equal to 0 and the largest value equal to 1. Normalized values were then averaged and plotted with standard the deviation. As individual growth traces varied on the timescale of a few minutes, some complementary montage examples may include timepoints outside of the plotted data range. However, plotted averages represent data normalized to the maximum time shown. Initial mean intensity measurements were derived from traces of growth events and defined as the intensity of the first frame of background subtracted punctate EGFP signal. Measurement of EGFP-EPS8 puncta intensities was performed in Nikon Elements using z-projected deconvolved images described above. In a cell shaped ROI, puncta were detected using the “Bright Spots” function and the 16-bit mean intensities were measured for the first frame of each time-lapse movie. Histograms of 16-bit mean intensities were generated using Prism V.7 (Graphpad). The microvillus growth rate was measured using maximum intensity projected images in FIJI. The length of the microvillus was measured at the first frame of detectable coalesced mCherry-ESPIN signal (Length_initial_) and then either the last frame of the movie, or the last frame with the microvillus mostly parallel to the cell surface (Length_final_). Events were excluded if the microvillus was mostly perpendicular to the cell surface. The growth rate was then calculated using (Length_final_ – Length_initial_)/(Time_final_-Time_initial_), with individual growth rates represented in the plot. The growth rate was calculated directly from growth events measured in normalized intensity plots, unless indicated in the figure legends.To measure the tip:cytosol ratio, intensity measurements were performed in FIJI using SIM images reconstructed using Nikon NSIM software. The EGFP or AlexaFluor488 signal was manually thresholded in each cell, and a rectangular ROI was drawn to measure the mean intensity at the tips of microvilli and in the cytoplasm, excluding the nuclear region. The same ROI area was used within a single cellular measurement, but not necessarily between cells, as to account for variation in cell size and shape. Measurement of microvilli collapse events was performed manually in FIJI. All measurements were performed on z-projected images deconvolved using the Richardson-Lucy algorithm in Nikon Elements software as described above. For all intensity and length measurements in all conditions, t=0 is defined as −10 frames before the characteristic drop in EGFP intensity (EGFP-EPS8, EGFP-IRTKS or Ezrin-GFP). A circular ROI was drawn to track the EGFP signal at the distal tips of microvilli. This same ROI was then used to measure CellMask-DeepRed intensity, when applicable. Circular ROIs of approximately the same size were used between independent collapse events. To measure microvilli lengths, a linear ROI was drawn to measure the change in microvilli length over time. Mean intensity values and lengths were first normalized and then averaged using Prism V.7 (GraphPad), and plotted with the standard deviation. As individual collapse intensity and length traces varied on the timescale of a few minutes, some complementary montage examples may include timepoints outside of the plotted data range. However, plotted averages represent data normalized to the maximum time shown.

### Statistical Analysis

Statistical significance was evaluated in Prism V.7 (GraphPad), by first running a D’Agostino and Pearson test to determine normality, and then an unpaired t-test for pairwise comparison (Supp. Fig 2G) or ANOVA with Kruskal-Wallis to compare more than two conditions (Supp. Figs 3 CE). For all figures, error bars represent standard deviation.

## Notes

### Competing Interest Statement

The authors have declared no competing interest.

