## Supplementary figures and images for "Direct visualization of epithelial microvilli biogenesis"

### Supplemental Figures 1-4

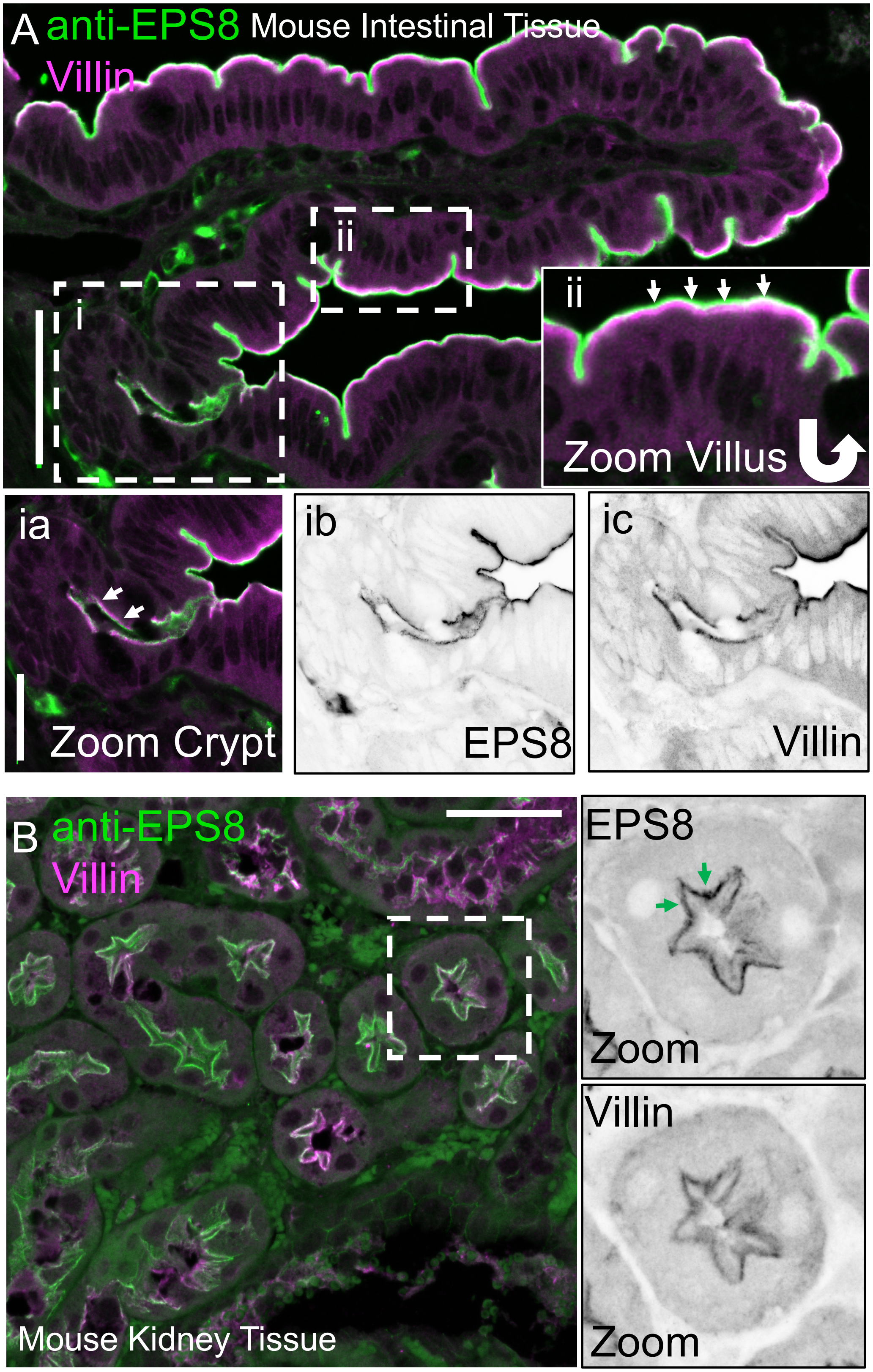

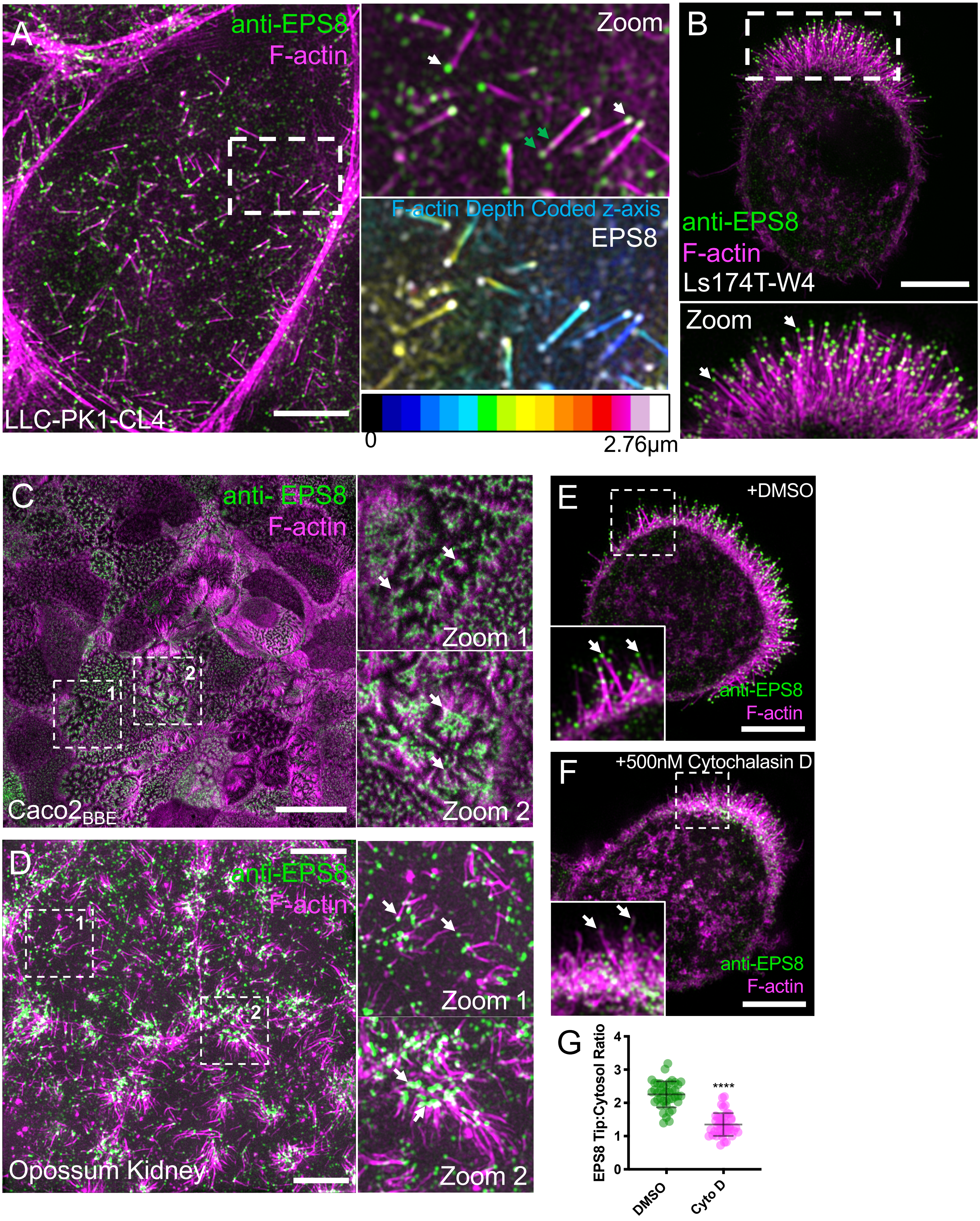

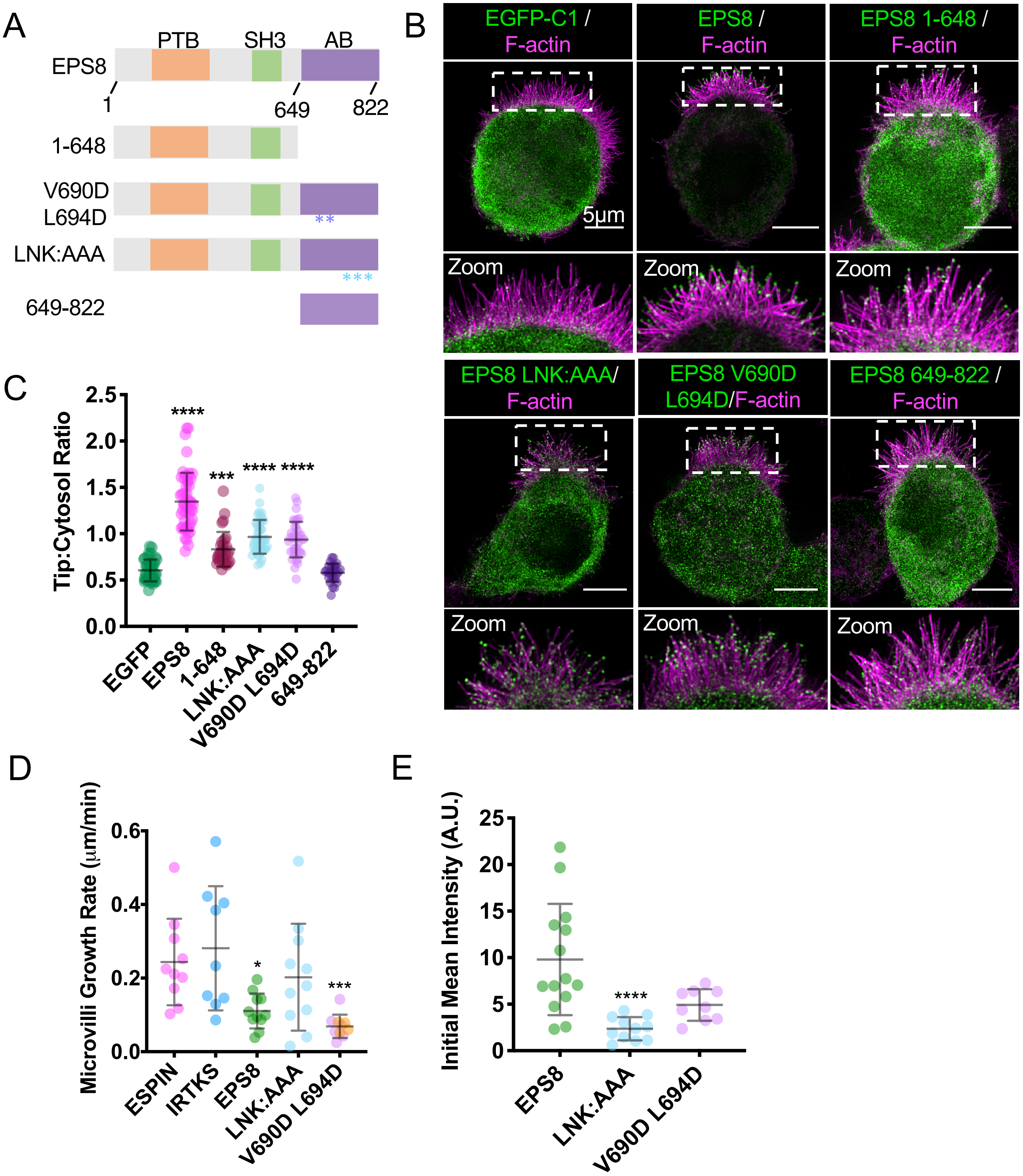

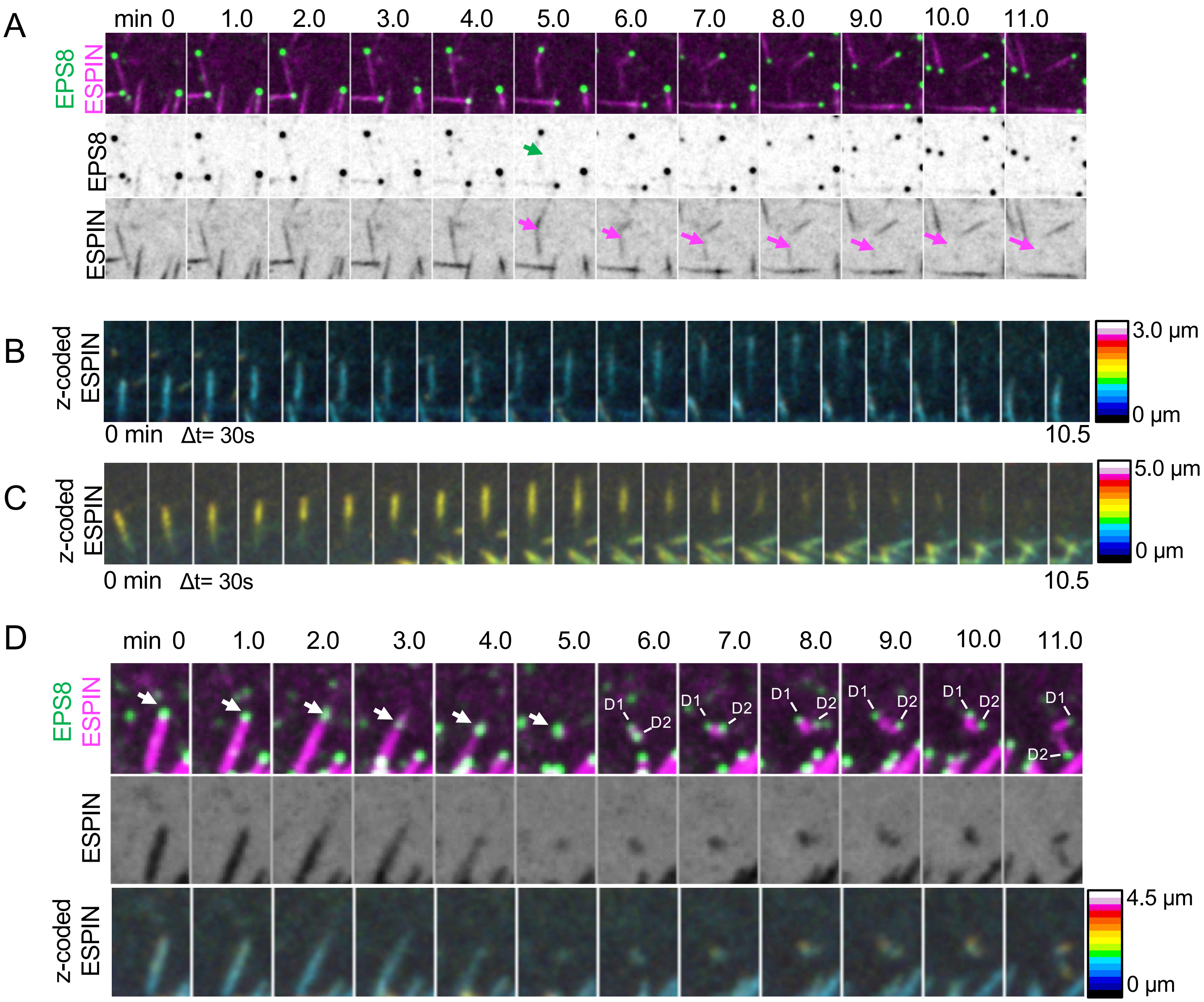
