## Supplemental Figure and Video Legends for "Direct visualization of epithelial microvilli biogenesis"

**SUPPLEMENTARY INFORMATION**

**FIGURE LEGENDS**

**Supplementary Figure 1**

(A) Representative image of paraffin embedded mouse intestinal tissue stained for endogenous EPS8 and the microvilli actin bundle marker villin. Scale bar = 50μm. Zoom boxes from i indicate the crypt compartment and ii indicates the villus. Arrows in ii and ia denote tip localization of EPS8. Scale bar for ia= 25μm. Images in A are from a single optical section acquired using confocal microscopy. (B) Representative images of paraffin embedded mouse kidney tissue stained for endogenous EPS8 and villin. Dashed box indicates zoom. Green arrows indicate EPS8 enrichment at the apical domain of kidney tubules. Scale bar = 50μm. Images in C are maximum intensity projections acquired using confocal microscopy.

**Supplementary Figure 2**

(A) Representative CL4 cell from an early stage of differentiation stained with an anti-EPS8 antibody and phalloidin to mark F-actin. Scale bar = 5μm. Zoom panels highlight individual microvilli and depth coding in the z plane. White arrows denote tip targeted EPS8. Occasionally, we see EPS8 localize along the side of the microvillus actin bundle (green arrows). (B) Representative image of W4 cell stained with an anti-EPS8 antibody and phalloidin. Arrows in zoom panel highlight EPS8 at the tips of microvilli. Scale bar = 5μm. (C) $Caco2_{BBE}$ cells at day 3 post confluency stained for EPS8 and phalloidin. Scale bar = 25μm. Zoom panels highlight EPS8 at the tips of microvilli. (D) Opossum kidney (OK) cells at day 2 post confluency stained for EPS8 and phalloidin. Scale bar = 5μm. Zoom panels highlight EPS8 at the tips of microvilli. (E) W4 cell treated with DMSO and stained for with an anti-EPS8 antibody and phalloidin. Dashed box indicates zoom, and arrows highlight EPS8 at the tips of microvilli. (F) W4 cell treated with 500nM cytochalasin D for 30 minutes and stained with an anti-EPS8 antibody and phalloidin. Dashed box indicates zoom and arrows highlight the de-enrichment of EPS8 from tips of microvilli. (G) Quantification of EPS8 tip enrichment for cells represented in E&F. ****$p<0.0001$ using an unpaired t-test. n=41 DMSO treated cells and n =43 cytochalasin D treated cells from 3 independent experiments. Images in A,B,D,E&F were acquired using structured illumination microscopy, image in C was acquired using confocal microscopy. All images are maximum intensity z projections.

**Supplementary Figure 3**

(A) EPS8 domain diagram. PTB=Phosphotyrosine Binding Domain, SH3 = Src-Homology 3, AB= Actin Binding. Putative capping and bundling point defective mutations are denoted by asterisks. (B) Representative images of EGFP only control and EGFP-EPS8 overexpression mutants in W4 cells imaged by SIM, displayed as maximum intensity projections. Dashed boxes indicate zoom and highlight protein localization in microvilli. F-actin is visualized by phalloidin. Images are not matched for brightness and contrast to show precise localization detail. (C) Quantification of tip to cytosol ratios of cells shown in B. Using ANOVA with Kruskal-Wallis, ****$p<0.0001$, ***$p=0.0008$, compared to EGFP only control. EGFP n= 41 cells; EPS8 n =49 cells; 1-648 n = 30 cells; V690D L694D n= 40 cells; LNK:AAA n=43 cells; 649-822 n=27 cells. Cells from all conditions are from least three independent transfections. (D) Quantification of microvillus growth rate. Using ANOVA with Kruskal-Wallis, *$p=0.0425$, ***$p=0.0002$ compared to cells expressing mCherry-ESPIN (ESPIN) only. ESPIN n=10 growth events, IRTKS n=9 growth events, EPS8 n=11 growth events, LNK:AAA n=11 growth events, V690D L694D n= 10 growth events. Each data point represents a single *de novo* microvillus growth event, corresponding to data shown in Figure1 D&E (EPS8), Figure2 B&D (LNK:AAA), and Figure2 C&E (V690D L694D). Data points highlighted in orange represent events that did not fit criteria to be included in intensity growth plots (Fig 2D). (E) Initial Mean Intensity values, signifying background subtracted 16-bit intensity from first frame of detectable EGFP signal for each condition. Each data point represents a single *de novo* microvillus growth event, corresponding to data shown in Figure1 D&E (EPS8), Figure2 B&D (LNK:AAA), and Figure2 C&E (V690D L694D). Using ANOVA with Kruskal-Wallis, ****$p<0.0001$ compared to EPS8. EPS8 n=14, LNK:AAA n=11, and V690D L694D n= 9 growth events.

**Supplementary Figure 4**

A) Representative montage of a microvillus bundle breaking, and the proximal end undergoing collapse (magenta arrows) in a CL4 cell expressing EGFP-EPS8 and mCherry-ESPIN. Note the lack of a significant concentration of EPS8 at the tip of the collapsing bundle (green arrow). Box width = 6.6 µm. (B) Z-coding of the mCherry-ESPIN channel in a CL4 cell also expressing EGFP-IRTKS and stained with CellMask-DeepRed, demonstrating the bundle does not drift out of frame. Related to Fig. 7A. Box width = 2.5µm. (C) Z-coding of the mCherry-ESPIN channel in a CL4 also expressing EGFP-Ezrin and stained with CellMask-DeepRed, demonstrating the bundle does not drift out of frame. Related to Fig. 7B. Box width = 2.5µm.

(D) Representative montage of a CL4 cell expressing EGFP-EPS8 and mCherry-ESPIN wherein a microvillus undergoes collapse, and new daughter microvilli grow from remnants of EPS8 puncta and Espin. Arrows denote EPS8 at the tip of the collapsing bundle, while D1 and D2 denote EPS8 at the tips of daughter bundles 1 and 2. Z-coded ESPIN is included to show the core bundle does not drift out of plane. Box width = 3µm.

**Video S1** Dynamic microvilli on the surface of a CL4 cell expressing EGFP-EPS8 and mCherry-ESPIN. Left: EGFP-EPS8 in green and mCherry-ESPIN in magenta. Right: EGFP-EPS8 in white and mCherry-ESPIN depth coded in z. Related to Figure 1A. Scale bar = 5μm.

**Video S2** *De novo* microvillus growth event in a CL4 cell expressing EGFP-EPS8 (green) and mCherry-Espin (magenta), related to Figure 1D and E. Scale bar = 1μm.

**Video S3** *De novo* microvillus growth event in a CL4 cell expressing EGFP-EPS8 (green) and mCherry-β-actin (magenta), related to Figure 2H and I. Scale bar = 1μm.

**Video S4** 3-dimensional rendering of a microvillus actin bundle (mCherry-Espin), demonstrating daughter microvilli are derived from the mother microvillus. The rendering is rotated so the daughter bundles are growing to the right of the mother bundle, related to Figure 3C.

**Video S5** *De novo* microvillus growth event in a CL4 cell expressing EGFP-IRTKS (cyan) and mCherry-Espin (magenta), related to Figure 4B and D. Scale bar = 1μm.

**Video S6** *De novo* microvillus growth event in a CL4 cell stained with CellMask-DeepRed (yellow), and expressing EGFP-IRTKS (cyan) and mCherry-Espin (magenta), related to Figure 5A and D. Scale bar = 1μm.

**Video S7** *De novo* microvillus growth event in a CL4 cell expressing Ezrin-GFP (green) and mCherry-Espin (magenta), related to Figure 5C and E. Scale bar = 1μm.

**Video S8** A microvillus collapse event in a CL4 cell expressing EGFP-EPS8 (green) and mCherry-Espin (magenta), related to Figure 6A and C. Scale bar = 1μm.

**Video S9** A microvillus collapse event in a CL4 cell stained with CellMask-DeepRed (yellow) and expressing EGFP-IRTKS (cyan) and mCherry-Espin (magenta), related to Figure 7A and C. Scale bar = 1μm.

**Video S10** A microvillus collapse event in a CL4 cell stained with CellMask-DeepRed (yellow) expressing Ezrin-GFP (cyan) and mCherry-Espin (magenta), related to Figure 7B and D. Scale bar = 1μm.
